# Achieving better than 3 Å resolution by single particle cryo-EM at 200 keV

**DOI:** 10.1101/141994

**Authors:** Mark A. Herzik, Mengyu Wu, Gabriel C. Lander

## Abstract

Technical and methodological advances in single-particle cryo-electron microscopy (cryo-EM) have expanded the technique into a resolution regime that was previously only attainable by X-ray crystallography. Although single-particle cryo-EM has proven to be a useful technique for determining the structures of biomedically relevant molecules at near-atomic resolution, nearly 98% of the structures resolved to better than 4 Å resolution have been determined using 300 keV transmission electron microscopes (TEMs). We demonstrate that it is possible to obtain cryo-EM reconstructions of macromolecular complexes at a range of sizes to better than 3 Å resolution using a 200 keV TEM. These structures are of sufficient quality to unambiguously assign amino acid rotameric conformations and identify ordered water molecules, features previously thought only to be resolvable using TEMs operating at 300 keV.

## Introduction

Technical breakthroughs in the field of three-dimensional single-particle cryo-electron microscopy (cryo-EM) have led to a period of explosive growth (1). Most notable has been the development of the direct electron detector (DED), which enables the direct counting of incident electrons, thereby improving the detective quantum efficiency (DQE) across spatial frequencies compared with traditional charge-coupled device (CCD) cameras (2). Moreover, the expedited readout rates of DED cameras allow for the collection of movies comprising multiple frames, effectively dose-fractionating each image over a standard exposure time. Movie collection also facilitates improved correction of beam-induced motion and/or stage drift through frame alignment, for which a multitude of programs are available (2–7). Such landmark developments, coupled with advancements in automated data collection, improved algorithms offered by processing packages (e.g. CRYOSPARC (8), EMAN2 (9), FREALIGN (10), RELION (11), SPARX (12), etc), and implementation of more powerful computational resources have led to visualization of biological macromolecular assemblies in near-native states at unprecedented resolutions (6, 13, 14).

However, these aforementioned advances have brought about new challenges in sample preparation, data acquisition, automation, training, and validation. Ongoing developments in hardware, software, and methodologies are attempting to address some of these problems with the ultimate goal of making cryo-EM readily accessible to scientists from broad backgrounds while expanding the capabilities of the technique to previously intractable and dynamic macromolecules.

To date, the vast majority of high-resolution cryo-EM reconstructions have been obtained using TEMs operating at 300 keV, such as the FEI Titan Krios or JEOL 3200FS(C), equipped with a direct electron detector. Microscopes operating at 300 keV offer minimized inelastic scattering and specimen charging over those at lower voltages, the benefits of which confer an additional advantage in imaging thicker biological specimens (15). As such, the pairing of a 300 keV TEM with a DED currently represents the ideal imaging setup for obtaining structures of biological macromolecules at near-atomic resolutions. However, the high cost of purchasing and operating high-end TEMs renders such a setup unfeasible for many institutions. Campbell *et al*. have previously reported the achievable resolution limit of an FEI TF20 TEM operating at 200 keV coupled with a DED to be up to ~3.7 Å (16). Recently, Li *et al*. have determined the structure of a cypovirus capsid to ~3.3 Å using a Talos Arctica TEM (200 keV) with a DED, however, this molecule benefits from high molecular weight and very high internal symmetry (17). Here we demonstrate that a 200 keV FEI Talos Arctica paired with a Gatan K2 Summit can produce cryo-EM reconstructions of *Thermoplasma acidophilum* 20S proteasome and rabbit muscle aldolase to ~3.1 Å and ~2.6 Å resolution, respectively. The quality of the resulting maps are comparable to similarly resolved reconstructions obtained using a 300 keV TEM, enabling confident identification of ordered water molecules and unambiguous assignment of amino acid rotameric conformations. Our work represents an unprecedented achievement in the resolution limit for structures obtained on a 200 keV TEM, proving that such TEMs are capable of reconstructing macromolecules of varying sizes and symmetries at resolutions previously ascribed only to 300 keV microscopes. We therefore propose that a 200 keV TEM such as the Talos Arctica, equipped with a direct detector is well suited for producing high-resolution reconstructions.

## Results

### Data Collection and Processing Strategy

Data were acquired using a base-model Talos Arctica TEM (i.e. no phase plate, spherical aberration corrector, or energy filter) operating in NanoProbe mode with an extraction voltage of 4150 V, a gun lens setting of 8, a spot size of 3 or 4, a C2 aperture size of 70 *μ*m, and an objective aperture size of 100 *μ*m. Careful alignment of the TEM was performed before each data collection to maximize parallel illumination (18) (**SI Figure 1**) and ensure Thon rings were visible beyond ~3 Å resolution in the power spectrum of aligned images collected over amorphous carbon. Coma-free alignment was performed according to Glaeser *et al*. (18) immediately prior to and daily during data collection, as implemented in Leginon (19). Data were collected automatically using Leginon and all image pre-processing was performed using the Appion pipeline (20). Images were acquired at a nominal magnification of 45,000× (calibrated pixel size of 0.91 Å at the detector level) with the Gatan K2 Summit DED operating in superresolution mode. To minimize the effects of beam induced motions during acquisition, samples were prepared on gold grids (21). Exposures were acquired at the center of holes using a beam that was large enough to cover the entirety of the hole and contact the surrounding gold substrate. We strived to achieve a particle concentration that maximized the number of particles contained within the holes without resulting in overlapping particles and/or aggregation. Each 20S proteasome movie was acquired over 17 seconds with a frame rate of 4 frames/sec and a dose rate of ~3.8 electrons/pixel/sec. Each aldolase movie was acquired over 11 seconds with a frame rate of 4 frames/sec and an average dose rate of ~5.1 electrons/pixel/sec. The total cumulative dose for all datasets was in the range of 65-68 electrons/Å^2^ (see **Methods**).

**Figure 1.**
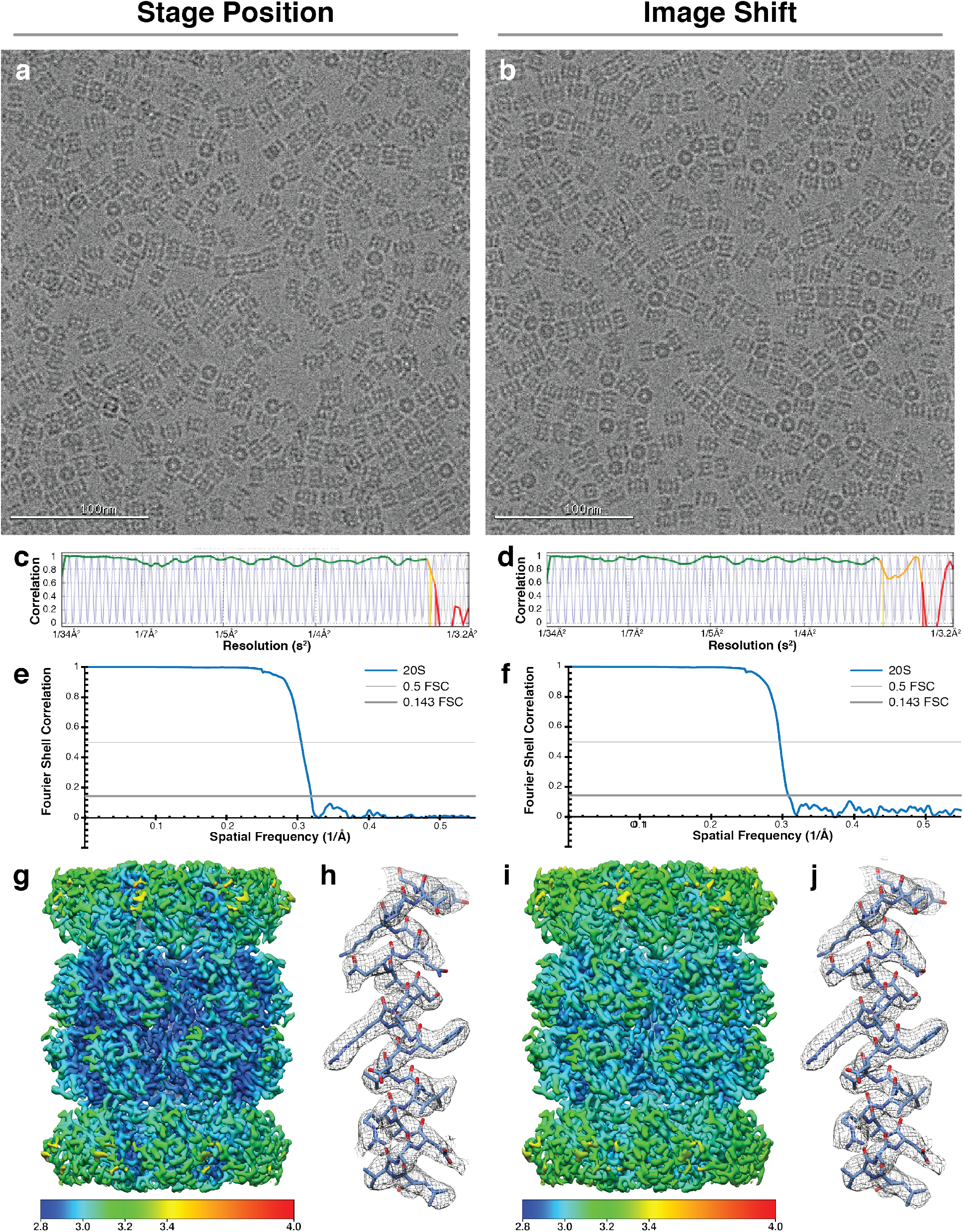
Cryo-EM reconstructions of the 20S proteasome at ~3.1 Å resolution. Motion-corrected micrographs of the *Thermoplasma acidophilum* 20S proteasome in vitreous ice acquired on a Talos Arctica TEM using either stage position (**a**) or image shift (**b**). Both micrographs were acquired at ~2 *μ*m underfocus. (**c** and **d**) CTF estimation fit (shown in blue) to experimental data and colored by CC (green, CC ≥90). Corresponding CTF estimates exhibit good fits to ~3 Å. Gold-standard Fourier shell correlation curves indicate a final resolution of 3.1 Å for the stage position dataset (**e**) and ~3.3 Å for the image shift dataset (**f**). Local resolution estimates of the final reconstructions calculated using BSOFT (30) reveal that the core of the molecule is resolved to better than 3 Å for both stage position (**g**) and image shift (**i**) reconstructions. An α-helix from the β-subunit (shown in stick representation) from each reconstruction with corresponding EM density (gray mesh, zoned 2 Å within atoms) exhibit clearly resolved side-chain density (**h** and **j**).

Mechanical and beam-induced motions were corrected and dose weighting performed using MotionCor2 (5) using a 5×5 patch size and a B-factor of 100. Whole-image contrast transfer function (CTF) estimation was performed using CTFFind4 (22) and micrographs yielding CTF estimates below an Appion confidence value of 0.9 were eliminated from further processing. Per-particle CTF estimates calculated using gCTF (23) were utilized for all subsequent processing steps. Reference-free 2D classification and subsequent 3D classification and 3D auto-refinement were performed using RELION 2.0 (24). All reported resolutions are based on the gold-standard criterion (25, 26) with all Fourier shell correlation (FSC) curves corrected for the effects of soft-masking by high-resolution noise substitution (27) (see **Methods**).

### Structures of the 20S proteasome at ~3.1 Å and ~3.3 Å resolution

In order to assess the resolution capabilities of our Talos Arctica TEM we sought to determine the structure of the 20S proteasome from *Thermoplasma acidophilum*, a symmetric ~700 kDa bacterial protein complex with a high thermal stability (thermoinactivation ~97°C (28)) that had previously been determined to ~2.8 Å resolution using cryo-EM (29). Briefly, 3 *μ*L of 1.5 mg/mL of purified 20S proteasome (hereafter referred to as 20S) was applied to Quantifoil UltrAuFoil^®^ grids containing 1.2 *μ*m holes spaced 1.3 *μ*m apart and had been plasma-cleaned in a Gatan Solarus plasma cleaner using a 75% argon / 25% oxygen atmosphere at 15 Watts for 6 seconds. Excess liquid was manually blotted for ~6 seconds at 4°C before immediately being plunged into liquid ethane cooled by liquid nitrogen (see **Methods**).

We collected 629 movies of frozen-hydrated 20S with a defocus range of −0.8 *μ*m to −1.8 *μ*m using stage position to navigate to each desired exposure target. The stage was allowed to settle for 30 seconds prior to each exposure in order to minimize the effects of stage movement-induced motion. Thon rings in the FFT of motion-corrected movies were visible to ~3 Å (**Figure 1**). 153,429 particles were extracted from the dose-weighted, aligned micrographs for reference-free 2D classification using RELION. 106,581 particles were then refined using the gold-standard protocol implemented in RELION to yield a map of ~3.3 Å resolution. Performing per-particle CTF estimation using gCTF (23) further improved the resolution of the map to ~3.1 Å. Local resolution estimation of this EM density using BSOFT (30) indicates that most of the map resolved to better than ~3.2 Å resolution, with the core of the molecule resolved to ~2.8 Å resolution (**Figure 1 and SI Figure 2**). Indeed, an overlay of the ~2.8 Å 20S proteasome reconstruction EMD-6287 reveals that the EM densities are of comparable quality, with both maps possessing clearly resolvable side-chain densities (**SI Figure 3**).

**Figure 2.**
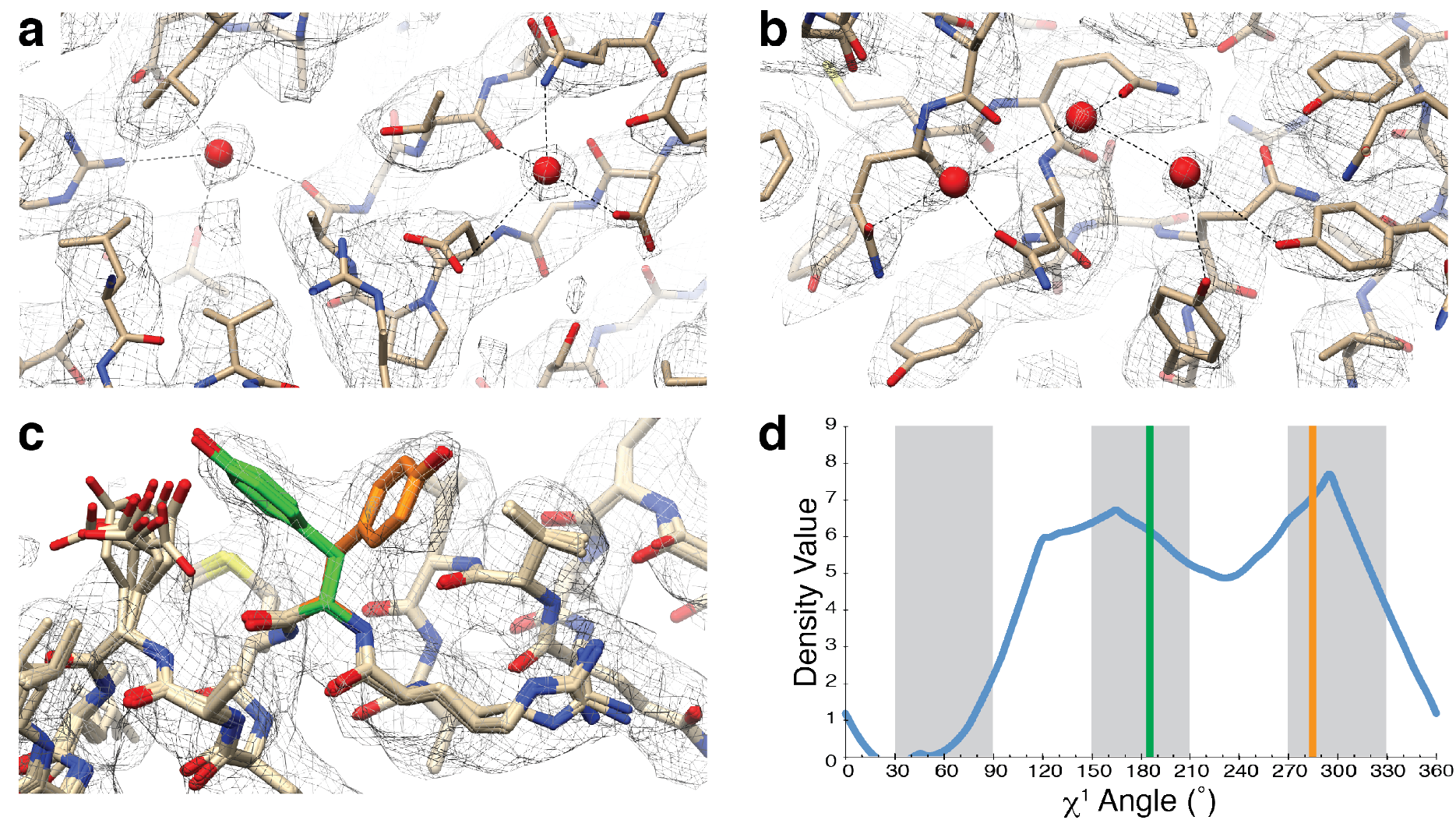
Resolving ordered water molecules and alternate rotameric conformations in the ~3 Å 20S proteasome reconstruction. (**a** and **b**) The ~3 Å 20S EM density (gray mesh) is of sufficiently high resolution to observe ordered water molecules (shown as red spheres). Hydrogen bonds to the water molecules are shown as black dotted lines. (**c**) The EM density clearly shows that Tyr58 of the β-subunit adopts two alternate rotameric positions. The top 10 models refined against the density distribute into these rotameric positions, colored either orange (60%) or green (40%). (**d**) EMRinger (55) analysis of the EM density corresponding to Tyr58 (blue line) confirms that the alternate conformations lie at ideal rotameric positions (thick gray bars). The refined χ^1^ angles for Tyr58 are shown as vertical lines and colored according to panel C (conformation 1 is colored orange and conformation 2 is colored green).

**Figure 3.**
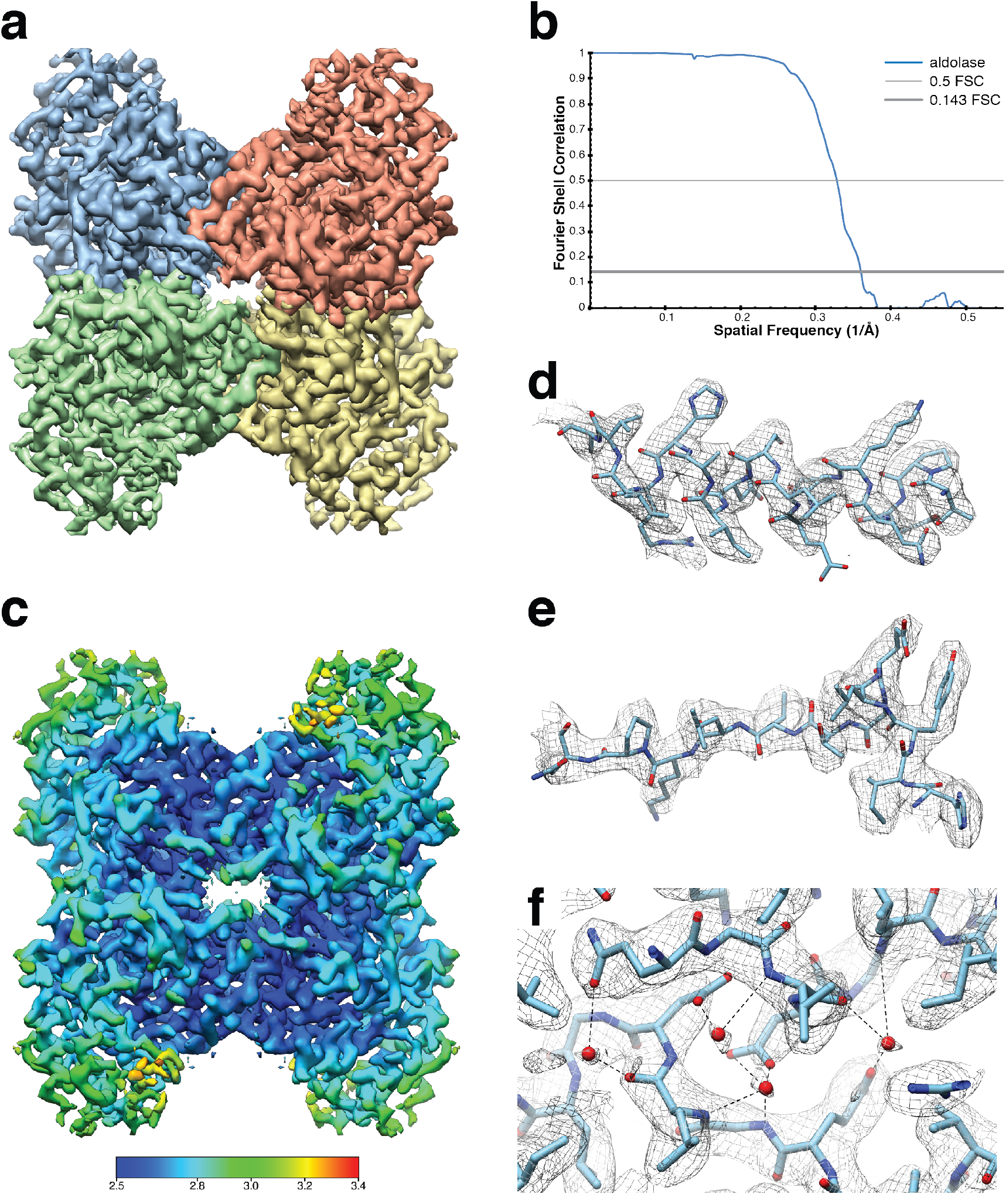
Structure of rabbit muscle aldolase at ~2.6 Å resolution. (**a**) The ~2.6 Å resolution aldolase EM density (D2 symmetric) segmented based on protomer organization. (**b**) The gold-standard Fourier shell correlation curve indicates a final resolution of 2.6 Å at 0.143 FSC. (**c**) The local resolution estimate of the final aldolase reconstruction calculated using BSOFT reveals that most of the molecule is resolved to better than 2.8 Å, with the core of the molecule resolved to ~2.5 Å. (**d and e**) Representative regions of the aldolase EM density (gray mesh, zoned 2 Å within atoms) indicate the map is of sufficient quality to unambiguously assign side-chain conformations as well as (**f**) the placement of putative ordered water molecules.

As an alternative data collection strategy to using stage position for navigating to each exposure target, we also collected a data set using stage position to first navigate to the center of 4 holes before image shifting the beam to each exposure target (**SI Figure 4**). This data collection strategy has the benefit of being able to collect more micrographs; on average, 18 images were collected per hour using image shift compared with 16 images per hour using stage position navigation (**Table 1**). However, substantial image shifting can introduce beam aberrations and decrease phase coherence. To determine the extent by which the loss of phase coherence due to beam tilting affects the attainable resolution, we collected 394 movies of frozen-hydrated 20S using image shift to navigate to acquisition targets. In order to eliminate as many non optically-related discrepancies as possible between the stage movement and image shift collections, all the data were collected using sample grids prepared and frozen in parallel during back-to-back sessions on the Talos Arctica TEM with similar acquisition parameters (i.e. magnification, dose rate, cumulative dose, etc.). Using a similar data processing and refinement strategy as employed for the stage position dataset, a final stack of 96,254 particles yielded a ~3.3 Å resolution reconstruction (**Figure 1 and SI Figure 5**). An estimation of the local resolution of this EM density indicates that most of the map resolves to ~3.2 Å resolution or better, with parts of the core resolved to better than 3 Å (**Figure 1**).

**Table 1.** Data collection, reconstruction, and model refinement statistics

|  | 20S Proteasome |  | Aldolase |
| --- | --- | --- | --- |
| **Data collection** |  |  |  |
| Microscope | Talos Arctica | Talos Arctica | Talos Arctica |
| Voltage (keV) | 200 | 200 | 200 |
| Nominal magnification* | 45,000x | 45,000x | 45,000x |
| Exposure navigation | Stage Position | Image Shift | Stage Position |
| Electron dose (e<sup>-</sup> Å<sup>-2</sup>) | 65 | 65 | 68 |
| Dose rate (e<sup>-</sup>/pixel/sec) | 3.8 | 3.8 | 5.1 |
| Detector | K2 Summit | K2 Summit | K2 Summit |
| Pixel size (Å)* | 0.92 (0.46) | 0.92 (0.46) | 0.92 (0.46) |
| Defocus range (μm) | -0.8 to -2.8 | -0.8 to -2.8 | -0.6 to -2.7 |
| Micrographs Collected (per hour) | ~16 | ~18 |  |
| Micrographs Used | 359 | 283 | 659 |
| Total extracted particles (no.) | 153,429 | 111,184 | 1,009,341 |
| Refined particles (no.) | 106,581 | 96,254 | 606,278 |
| **Reconstruction** |  |  |  |
| Final particles (no.) | 106,581 | 96,254 | 83,910 |
| Symmetry imposed | D7 | D7 | D2 |
| Resolution (global) | 3.14 Å | 3.28 Å | 2.6 Å |
| FSC 0.5 (unmasked/masked) | 3.73/3.28 Å | 3.79/3.37 Å | 3.76/2.84 Å |
| FSC 0.143 (unmasked/masked) | 3.37/3.14 Å | 3.45/3.26 Å | 3.24/2.62 Å |
| Applied B-factor (Å<sup>2</sup>) | -101 | -99 | -35 |
| **Refinement\*\*** |  |  |  |
| Protein residues | 5936 | 5936 | 1372 |
| Ligands/waters (atoms) | 560 | 420 | 80 |
| Map Correlation Coefficient |  |  |  |
| Global | 0.61 (±0.01) | 0.60 (±0.01) | 0.64 (±0.01) |
| Local | 0.81 (±0.01) | 0.79 (±0.01) | 0.87 (±0.01) |
| R.m.s. deviations |  |  |  |
| Bond lengths (Å) | 0.009 (±0.001) | 0.010 (±0.001) | 0.011 (±0.001) |
| Bond angles (°) | 1.024 (±0.071) | 1.040 (±0.064) | 0.839 (±0.023) |
| Ramachandran |  |  |  |
| Outliers | 0.10 (±0.12) | 0.07 (±0.16) | 0.06 (±0.12) |
| Allowed | 3.50 (±0.55) | 4.37 (±0.91) | 2.76 (±0.49) |
| Favored | 96.40 (±0.53) | 95.56 (±0.91) | 97.18 (±0.48) |
| Poor rotamers (%) | 2.00 (±0.70) | 1.89 (±1.19) | 0.14 (±0.19) |
| MolProbity score | 1.63 (±0.16) | 1.65 (±0.16) | 1.53 (±0.06) |
| Clashscore (all atoms) | 4.49 (±0.62) | 4.92 (±0.48) | 9.12 (±1.13) |
| Mean per-residue Cα RMSD (Å) | 0.37 | 0.44 | 0.21 |
| Per-residue Ca RMSD range (Å) | 0.04-1.15 | 0.05-2.50 | 0.01-1.39 |
| EMRinger Score(55) | 4.14 (±0.47) | 4.05 (±0.29) | 4.49 (±0.23) |

\*Calibrated pixel size at the detector
\*\*Values in parentheses correspond to values calculated from top 10 models

Comparison of the two 20S reconstructions generated using our 200 keV TEM reveals that maps of comparable quality can be obtained with less than a ~0.2 Å resolution loss (nominal, gold-standard FSC) when using image shift versus stage position for exposure target navigation. Visual inspection of the 20S EM densities indicates only modest differences in side-chain resolvability (**Figure 1**). This result suggests that the extent of the loss of phase coherence associated with the magnitude of beam tilt used in our image shift navigation strategy minimally disrupts optimal image formation, with marginal loss of EM density quality. However, image shifting substantially beyond the 1.76 *μ*m distance used for this study will likely deteriorate the phase coherence and attainable resolution.

The final 20S reconstruction from the stage movement data set features visible side-chain density for most of the molecule with the best-resolved regions of the EM map possessing density that correspond to numerous well-ordered water molecules (**Figure 2**). Importantly, the location of many of these ordered waters correlate with those previously observed for the *T. acidophilum* 20S proteasome determined by cryo-EM (EMD-6287 (29)) and X-ray crystallography (PDB ID: 1YAR (31)), corroborating the placement of these waters (**SI Figure 3**). Further examination of the density reveals that some well-ordered side chains exhibit alternate conformations that lie within expected rotameric positions (**Figure 2**). To the best of our knowledge, these are the first observations of water molecules and/or alternate side-chain conformations in a reconstruction obtained using a 200 keV TEM.

### Structure of rabbit muscle aldolase at ~2.6 Å resolution

Nearly all structures determined to better than 5 Å resolution using a 200 keV TEM possess high internal symmetry (EMDs-2791 (16); 5886 (32)) and/or comprise a molecular weight of ≥700 kDa (EMDs-5247 (33); 5592 (34); 6458 (35)). Only one example exists in the EMDB that does not fulfill either of these two criteria (EMDs-6337 (36)). In fact, only 1 structure less than 200 kDa has been reported to better than 3 Å resolution (EMD-8191 (13)), and it was determined using a Titan Krios TEM at 300 keV. In order to determine whether near-atomic resolution was feasible for complexes in this size range using a 200 keV TEM, we sought to determine the structure of aldolase, a small homotetrameric glycolytic enzyme with a molecular weight of ~150 kDa. Briefly, lyophilized aldolase purified from rabbit muscle (Sigma Aldrich) was resuspended in 20 mM HEPES pH 7.5, 50 mM NaCl and subjected to size-exclusion chromatography. The peak fractions corresponding to native aldolase, as determined by SDS-PAGE, were pooled and concentrated to ~1.6 mg/mL. 3 *μ*L of concentrated aldolase was applied to a plasma-cleaned Quantifoil UltrAuFoil^®^ grid with 1.2 *μ*m holes spaced 1.3 *μ*m apart, using the same plasma settings as the 20S. Excess liquid was manually blotted for ~4 seconds at 4°C before immediately plunge-freezing in liquid ethane cooled by liquid nitrogen (see **Methods**).

784 super-resolution movies of frozen-hydrated aldolase were collected using a nominal defocus range of −0.8 *μ*m to −1.4 *μ*m using stage position to navigate to each exposure target. Similar to acquisition settings used for the 20S proteasome, a stage settling time of 30 seconds was used prior to each exposure to dampen the effects of stage movement-induced motion. Thon rings in the FFT of motion-corrected aldolase movies were visible to ~3 Å. 1,009,341 particles were extracted from the aligned, radiation damage-corrected summed frames, decimated by a factor of 4, and subjected to reference-free 2D classification using RELION 2.0 (24). Those particles exhibiting secondary structure elements were selected for further refinement. Following several rounds of 3D refinement and classification, 83,910 refined particles yielded a ~2.6 Å resolution reconstruction as determined by gold-standard FSC (see **Methods, Figure 3**, and **SI Figure 5**) (25, 26). Local resolution estimates using BSOFT (30) reveals that most of the map is resolved to better than 2.8 Å with the most ordered regions of the map resolved to ~2.5 Å (**Figure 3**).

The final reconstruction possesses clear side-chain density for most of the molecule and well-resolved backbone density. The best-refined model agrees well with previously published structures determined by X-ray diffraction methods, with an overall RMSD of 0.45 Å (PDB ID: 6ALD) (37). Further examination of the density reveals that the best resolved regions of the map possess density attributed to ordered water molecules (**Figure 3**), as expected for an EM reconstruction determined to better than 3 Å resolution. Moreover, the positions of these water molecules are conserved between those identified in 6ALD (**SI Figure 7**). Together, these results exemplify the capabilities of 200 keV TEMs in resolving biomedically relevant macromolecules to near-atomic resolution.

## Discussion

Transmission electron microscopes (TEMs) operating at 300 keV have been shown to be optimal for imaging thicker biological specimens due to reduced inelastic scattering and smaller scattering angles, which reduces the beam spreading and image blurring that is associated with charging effects and radiolysis (15, 38). The deleterious effects of beam-induced motion and radiation damage have been remediated through fractionation into movie frames using a direct electron detector (DED) with subsequent image alignment and dose-weighting (2–7). For these reasons, 300 keV TEMs such as the FEI Titan Krios coupled with DEDs remain the flagship instruments in the field of high-resolution single-particle cryo-EM, accounting for nearly 98% of all reconstructions reported to 4 Å resolution or better. The results presented in this study demonstrate that cryo-EM reconstructions resolved to ~3 Å resolution and better are achievable using a 200 keV TEM coupled with a K2 Summit direct electron detector for samples of different sizes and internal symmetries. The maps generated herein are of sufficient quality for *de novo* model building as well as unambiguous identification of ordered waters and alternate side-chain conformations (**Figures 2 and 3**).

Several studies have used *T. acidophilum* 20S proteasome as a metric for characterizing the potential of TEMs and imaging accessories (2, 14, 16, 29, 39). We similarly sought to assess the resolution limit of the Talos Arctica using the 20S proteasome and two data collection strategies. We anticipated that a reconstruction obtained from images collected using image shift-based exposure targeting would be hindered by loss of phase coherence due to beam tilting. However, the difference in resolution between the two structures was minimal (<0.2 Å loss), with both data collection strategies yielding maps of comparable quality (**Figure 1**), indicating that the magnitude of beam tilt used for exposure targeting in this study (1.76 *μ*m) did not result in a substantial loss of phase coherence. Notably, both maps exhibited clear density attributed to ordered water molecules, an unprecedented achievement for structures obtained using a 200 keV TEM. The presence of alternate amino acid rotamers in the density map further attests to the quality of the reconstruction and indicates selective conformational switching within the protein. As such, we propose that using image shift navigation is a viable data collection strategy to maximize acquisition time, so long as one ensures parallel illumination of the beam and careful monitoring of excess defocus and astigmatism.

To contrast the 20S proteasome, which benefits from size, high internal symmetry, and high thermostability, and to assess the resolving capacity of a 200 keV TEM for smaller complexes, we also imaged rabbit muscle aldolase. Remarkably, using similar illumination conditions as those utilized for determining the structures of 20S proteasome, the final aldolase reconstruction achieved a resolution of ~2.6 Å (goldstandard FSC). In contrast, the resolution of the 20S reconstructions were limited to ~3 Å. While the smaller size of aldolase and thus the fewer voxels utilized in the calculation of gold-standard FSC may partly influence the higher reported resolution for aldolase, it is also likely that the smaller size of aldolase contributes to a more isotropic resolution that is not negatively influenced by flexible peripheral domains. However, differences in ice thickness and overall sample stability cannot be ruled out as contributors.

Approximately 20% of the 20S stage position reconstruction comprises particles imaged at greater than 2 *μ*m underfocus. Omitting these particles did not significantly alter the nominal resolution of the resulting reconstruction (**SI Figure 2**), indicating that these micrographs did not appreciably contribute structural information in the highest resolution frequencies. Furthermore, only 0.1% of the total particles contributing to the final aldolase reconstruction possessed defocus values greater than −1.5 *μ*m, again indicating that particles imaged at greater defoci made negligible contributions to highresolution information. These results corroborate the standard practice of maintaining the lowest defocus range necessary to produce low-resolution image contrast for particle alignment while limiting high-resolution dampening from the CTF envelope function (40). In order to maximize the envelope function for high-resolution data collection, it is essential to collect using minimal defocus values on a 200 keV TEM.

Typically, micrometer underfocus values have been prescribed for imaging targets less than 200 kDa (41). More recently, strategies implementing phase plates combined with a small amount of defocus to boost image contrast have been demonstrated to be an effective method for resolving small protein complexes (42). However, despite the small size of the aldolase particles used in this study, micrographs of sufficient contrast were obtained using comparable illumination conditions and similarly low underfocus values (as low as −0.6 *μ*m) as for the 20S proteasome, a comparatively larger and more structurally featureful target (Table 1). We believe this is largely attributed to the thinness of the ice of the aldolase specimen coupled with high particle density to boost particle contrast in the collected micrographs and facilitate alignment of movie frames. As such, our results suggest it is feasible to achieve near-atomic resolution reconstructions of a sub-200 kDa specimen using a conventional defocus approach, provided the particles are embedded in sufficiently thin ice.

Although we do not envision a 200 keV scope achieving reconstructions to better than ~2 Å resolution, we do anticipate that several experimental parameters can be further modified to improve the achievable resolution to beyond those obtained in this study. For instance, inclusion of a quantum energy filter would aid in eliminating some of the deleterious effects of inelastic scattering, which is particularly exacerbated at lower acceleration voltages (15). We also anticipate that using a smaller C2 aperture (i.e. 30 *μ*m) would further improve illumination of the sample by decreasing the effects of spherical aberration, the primary limitation to achieving the theoretical resolution for a TEM (43). Additionally, increasing the nominal magnification at specimen level would place imaging conditions at a higher camera DQE, thus yielding higher image signal-to-noise ratio and particle contrast (44), albeit at the expense of particle number and the ability to estimate the contrast transfer function due to the smaller field of view.

A major advantage of single-particle cryo-EM is the ability to determine the structures of biological specimens in near-native environments without the requirement of a crystal lattice. However, most structures that have been determined to near-atomic resolution have utilized a Titan Krios or JEOL 3200FS(C) TEM operating at 300 keV. Herein we examined the *T. acidophilum* 20S proteasome and rabbit muscle aldolase, as complementary metrics to show that a base-model 200 keV Arctica TEM equipped with a DED is also capable of near-atomic resolution cryo-EM reconstructions of biological macromolecules of varying sizes and symmetries. These specimens were selected for their robust structural integrity and limited conformational heterogeneity in order to probe the resolution capabilities of this instrumentation, and we recognize that not all samples are as well behaved as those presented in this study. Nonetheless, our findings serve as an important proof-of-principle for near-atomic resolution structure determination on a 200 keV TEM. Given that the 200keV Talos Arctica used in this study costs less than roughly half that of the 300keV Titan Krios, it is our hope that this work expands the field of high-resolution cryo-EM to a broader range of research institutes that are unable to incur the steep long-term financial commitment of a 300 keV TEM.

## Materials and Methods

### Cryo-electron microscopy sample handling and grid preparation

Pure aldolase isolated from rabbit muscle was purchased as a lyophilized powder (Sigma Aldrich) and solubilized in 20 mM HEPES (pH 7.5), 50 mM NaCl at ~3 mg/ml. Aldolase was further purified by size-exclusion chromatography using a Sepharose 6 10/300 (GE Healthcare) column equilibrated in solubilization buffer. Peak fractions were pooled and concentrated to 1.6 mg/ml immediately prior to cryo-electron microscropy (cryo-EM) grid preparation.

Archaeal 20S proteasome (*T. acidophilum*) was kindly donated by Drs. Zanlin Yu and Yifan Cheng at The University of California, San Francisco and used as is without further modification.

For cryo-EM, 3 μL of purified aldolase (1.6 mg/ml) or 20S proteasome (0.5 mg/ml) were dispensed on freshly plasma cleaned UltrAuFoil^®^ R1.2/1.3 300-mesh grids (Electron Microscopy Services) and manually blotted using filter paper (Whatman No.1) for ~4 seconds (≥95% relative humidity, 4°C) immediately prior to plunge freezing in liquid ethane cooled by liquid nitrogen.

### Cryo-EM data acquisition, image processing, and refinement

Microscope alignments were performed on a cross-grating calibration grid. Condenser alignments and stigmation were performed as described previously for a two condenser lens system (45). Proper eucentric height of the specimen was determined using Leginon prior to setting focus. Parallel illumination of the beam was maximized in diffraction mode by first adjusting the defocus to bring the objective aperture into focus in the front focal plane of the diffraction lens followed by adjustments of beam intensity to minimize the spread of powder diffraction (46). The resulting beam intensity value was saved in Leginon and remained unchanged throughout data collection. The objective aperture was centered and any objective lens astigmatism was minimized. Dose rate measurements on the K2 Summit DED were then collected to determine whether changes to spot size were necessary to achieve the desired dose rate. In the event that changes in dose rate were required, as was the case for increasing per frame cumulative dose during aldolase imaging, adjustments to spot size were made and alignments were repeated, including maximization of parallel illumination. Coma-free alignment was then performed as described previously (18). Daily adjustments, if necessary, were made during data collection to maintain lens stigmation and ensure the beam was coma-free.

All cryo-EM data were acquired using the Leginon automated data-acquisition program (19). All image pre-processing (frame alignment, CTF estimation, particle picking) were performed in real-time using the Appion image-processing pipeline (20) during data collection.

Images of frozen hydrated aldolase or 20S proteasome were collected on a Talos Arctica transmission electron microscope (TEM) (FEI) operating at 200 keV. Movies were collected using a K2 Summit direct electron detector (Gatan) operating in superresolution mode (super-resolution pixel size 0.46 Å/pixel) at a nominal magnification of 45,000x corresponding to a physical pixel size of 0.92 Å/pixel.

629 movies (68 frames/movie) of 20S proteasome were collected using stage position to navigate to the exposure target. Movies were collected over 17 seconds exposure with a dose rate of 3.8 e^−^/pix/s, resulting in a total dose of ~65 e^−^/Å^2^ (0.96 e^2^/Å^2^/frame) and a nominal defocus range from −0.8 *μ*m to −2.8 *μ*m. The same exposure settings were used to collect 394 movies using image shift target navigation. Superresolution images were Fourier-binned by 2 (0.92 Å/pix) prior to motion correction and dose-weighting using the MotionCor2 frame alignment program (5) as part of the Appion pre-processing workflow. Frame alignment was performed on 5 × 5 tiled frames with a B-factor of 100 applied. Unweighted summed images were used for CTF determination using CTFFIND4 (22). DoG picker (47) was used to automatically pick particles from the first 10 dose-weighted micrographs yielding a stack of 4,821 particles that were binned 8 × 8 (3.68 Å/pixel, 92 pixel box size) and subjected to reference-free 2D classification using an iterative topology-representing network-based classification followed by multireference alignment(48) in the Appion pipeline. The best 4 classes were then used for template-based particle picking against the each dataset using FindEM (49). Only micrographs fulfilling the following two criteria: 1) possessing a CTF estimate confidence of fit ≥90% and 2) with resolution estimates to 5 Å or better at CC=50% (359 total from the stage position data set; 283 total from the image shift data set) were used.

For the 20S proteasome data set collected using stage position, 153,429 particles were extracted rom the dose-weighted summed images, further binned 4 × 4 (3.68 Å/pix, 92 pixel box size), and subjected to reference-free 2D classification using RELION 2.0 (24). The particles from the best-aligned classes (106,581 particles) were then 3D auto-refined with D7 symmetry imposed using EMD-6283 as an initial model (low-passed filtered to 60 Å) to yield a ~7.4 Å resolution reconstruction (7.36 Å Nyquist). Particles were then re-centered using the refined particle offsets before being reextracted unbinned (0.92 Å/pix, 368 pixel box size) for 3D auto-refinement using a scaled output from the 4 × 4 binned refinement as an initial model. The refinement was continued using a soft-mask (3 pixel extension, 5 pixel soft cosine edge) generated from a volume contoured to display the full density. The final resolution was estimated to 3.32 Å (gold-standard Fourier shell correlation (FSC) (25, 26)) using phase-randomization to account for the convolution effects of a solvent mask on the FSC between the two independently refined half maps (27). The refined particle coordinates were then used for local CTF estimation using gCTF (23) followed by re-extraction of particles using a 512 pixel box size to limit aliasing in the Fourier domain. Gold-standard 3D autorefinement using the same soft mask yielded a 3.14 Å reconstruction (gold-standard FSC) (**Supplementary Figure 1**). These particles were then subjected to 3D classification (k=4, tau fudge=25) without angular or translational searches using the same soft mask. However, further refinement of each individual class did not yield an improvement in nominal FSC-reported resolution. In addition, further filtering of the final particle stack using a cutoffs based on the height of the probability distributions at their maximum (MaxValueProbDistribution) did not result in an increase in nominal FSC-reported resolution. Finally, removal of all particles with an estimated defocus of greater than 2 *μ*m yielded a reconstruction 0.01 Å lower in resolution indicating that particles illuminated at greater defoci did not significantly contribute high-resolution information (**Supplementary Figure 1**).

Processing and refinement of the 20S proteasome data set collected using image shift was performed similarly as described for the 20S proteasome data set collected using stage position.

810 movies (44 frames/movie) of rabbit muscle aldolase were acquired over 11 s with a dose rate of 5.1 e^−^/pix/s, yielding a total dose of ~68 e^−^/Å^2^ (1.55 e^−^/Å^2^/frame), and a nominal defocus range from −0.8 *μ*m to −1.4 *μ*m. 2 × 2 Fourier-binned super-resolution images (0.92 Å /pix) were motion correction and dose-weighted using the MotionCor2 program (5) using 5 × 5 tiled frames with a B-factor of 100 applied. Unweighted summed images were used for CTF determination using CTFFIND4 (22). Weighted sums were used for automated template-based particle picking with FindEM (49). Only those micrographs with CC ≥90% and with resolution estimates 5 Å or better at CC=50% (659 total) were used. 1,009,341 particles were extracted from these micrographs and binned 4 × 4 (3.68 Å/pix, 92 pixel box size). Reference-free 2D classification in RELION 2.0 (24) was then used to sort out non-particles and poor-quality picks in the data. A total of 606,278 particles corresponding to 2D class averages that displayed strong secondary-structural elements were selected for 3D auto-refinement with D2 symmetry imposed using a simulated EM density generated from the previously published X-ray crystal structure of rabbit muscle aldolase (PDB ID: 6ALD (37)) low-pass filtered to 30 Å. The refined particle coordinates were then used for local CTF estimation using gCTF (23) followed by re-extraction of particles binned 2 × 2 (1.84 Å/pix) followed by subsequent 3D auto-refinement using the 4 × 4 binned map as an initial model. The refinement was continued using a soft mask (5 pixel extension, 10 pixel soft cosine edge) generated from a volume contoured to display the full density. The final resolution was estimated to 3.64 Å (Nyquist=3.64 Å, gold-standard FSC (25, 26)) using phase-randomization to account for the convolution effects of a solvent mask on the FSC between the two independently refined half maps (27). These particles were then subjected to 3D classification (k=6, tau fudge=8) without angular or translational searches using the same soft mask. The classes that possessed the best resolved side-chain and backbone densities were re-centered and re-extracted (83,910 particles, 0.92 Å/pixel, 368 pixel box size). These particles were then 3D auto-refined using a soft mask. The final resolution was estimated to ~2.6 Å (gold-standard FSC) (**Supplementary Figure 1**).

### Model building and refinement

For each of the final reconstructions, an initial model was subjected to a multimodel pipeline using methodologies similar to those described previously (50). Briefly, for the 20S proteasome reconstructions, PDB ID:1YAR (31) was stripped of all ligands (i.e. PA26 and waters) and all alternate conformations, all occupancies were set to unity, all B-factors were set to a singular value, and all Ramachandran and geometric outliers were corrected. For aldolase, PDB ID: 6ALD was stripped of all cofactors and water molecules, with all occupancies set to zero and all Ramachandran and geometric outliers corrected. Initial docking of PDB ID: 6ALD into the density revealed additional density beyond the Cβ of alanine 146. Analysis of the UniProt (51) metadata associated with PDB ID: 6ALD (37) indicated this amino acid should be a lysine. Thus, lys146 was corrected in each chain prior to refinement. These initial models were then refined into the EM density using the imposed symmetry and estimated resolution while adjusting the Rosetta weighting and scoring functions according to the estimated map resolution.

Each of the 100 Rosetta-generated models (52) were ranked based on the number of Ramachandran outliers (%, lower better), geometry violations (%, lower better), Rosetta aggregate score (value, lower better), and MolProbity clashscore (53) (value, lower better). The top 10 structures that scored the best across all categories were selected for real-space refinement using the Phenix refinement package (54). Model–model agreement statistics were calculated using a previously described approach (50).

## Acknowledgements

We thank Jean-Christophe Ducom at The Scripps Research Institute High Performance Computing for computational support, and Bill Anderson at The Scripps Research Institute electron microscopy facility for microscope support. We are grateful to Yifan Cheng and Zanlin Yu for kindly providing the T20S sample used in this study. M.A.H. is supported by a Helen Hay Whitney Foundation postdoctoral fellowship. G.C.L. is supported as a Searle Scholar, a Pew Scholar, and by the National Institutes of Health (NIH) DP2EB020402. Computational analyses of EM data were performed using shared instrumentation funded by NIH S10OD021634.

## Data Deposition

The coordinates for the 20S stage position, 20S image shift, and aldolase structures are deposited in the Protein Data Bank with the PDB IDs 5VY3, 5VY4, and 5VY5, respectively. The corresponding EM density maps have been deposited to the EMDB with the IDs 8741, 8742, and 8743, respectively.

